# Ubiquitylation activates a peptidase that promotes cleavage and destabilization of its activating E3 ligases and diverse growth regulatory proteins to limit cell proliferation in Arabidopsis

**DOI:** 10.1101/090563

**Authors:** Hui Dong, Jack Dumenil, Fu-Hao Lu, Li Na, Hannes Vanhaeren, Christin Naumann, Maria Klecker, Rachel Prior, Caroline Smith, Neil McKenzie, Gerhard Saalbach, Liangliang Chen, Tian Xia, Nathalie Gonzalez, Mathilde Seguela, Dirk Inze, Nico Dissmeyer, Yunhai Li, Michael W. Bevan

**Author notes:** Co-first Authors (Hui Dong, Jack Dumenil, Fu-Hao Lu, Li Na). Co-Senior Authors (Yunhai Li, Michael W. Bevan).

## Abstract

The characteristic shapes and sizes of organs are established by cell proliferation patterns and final cell sizes, but the underlying molecular mechanisms coordinating these are poorly understood. Here we characterize a ubiquitin-activated peptidase called DA1 that limits the duration of cell proliferation during organ growth in *Arabidopsis thaliana*. The peptidase is activated by two RING E3 ligases, BB and DA2, which are subsequently cleaved by the activated peptidase and destabilized. In the case of BB, cleavage leads to destabilization by the RING E3 ligase PRT1 of the N-end rule pathway. DA1 peptidase activity also cleaves the de-ubiquitylase UBP15, which promotes cell proliferation, and the transcription factors TCP15 and TCP22, which promote cell proliferation proliferation and repress endoreduplication. We propose that DA1 peptidase activity regulates the duration of cell proliferation and the transition to endoreduplication and differentiation during organ formation in plants by coordinating the destabilization of regulatory proteins.

## INTRODUCTION

The shapes and sizes of organs are established by mechanisms that orient cell proliferation and determine the final numbers and sizes of cells forming the organ. Transplantation experiments showed that some animal organs have an intrinsic mechanism that determines their final size by controlling the duration of cell proliferation (Barry and Camargo 2013), which is controlled in part by the HIPPO/YAP pathway that limits cell proliferation and promotes apoptosis (Pan 2010). However, the mechanisms coordinating cell proliferation and cell size during organ growth remain poorly understood (Johnston and Gallant 2002). Due to the simpler planar structures of their organs such as leaves and petals, and the absence of cell movement due to rigid cell walls, plants have some experimental advantages for studying organ growth (Green et al. 2010).

Leaf growth in plants is initiated at shoot meristems (reviewed by Sluis and Hake 2015). After specification of boundaries and growth axes, the leaf lamina grows in an initial period of cell division in which cell size is relatively constant, followed by a transition to endoreduplication associated with cell expansion and differentiation (Breuer et al. 2010; De Veylder et al. 2011). The transition from cell proliferation to cell expansion is spatially and temporarily regulated during leaf growth, and appears to progress from the tip to the base of the leaf as a cell division arrest front (Kazama et al. 2010), accompanied by shifts in gene expression patterns (Efroni et al. 2008; Andriankaja et al. 2012). A key question is how the transition from cell proliferation to cell expansion and differentiation is coordinated to generate a correctly sized organ.

The RING E3 ligases Big Brother (BB) (Disch et al. 2006a) and DA2 (Xia et al. 2013) limit the duration of cell proliferation during organ growth. Members of the *DA1* family also limit cell proliferation (Li et al. 2008), and loss of function mutations in *BB* and *DA2* interact synergistically with the *da1-1* allele of *DA1* to increase organ and seed size in Arabidopsis (Li et al. 2008; Xia et al. 2013), suggesting one of their growth limiting activities is mediated by enhancing the growth-repressive activity of DA1 family members. Genetic analyses showed that DA1 reduced both the stability of UBP15 (Du et al., 2014), a de-ubiquitylation enzyme promoting cell proliferation (Lui et al., 2008), and reduced the stabilities of TEOSINTE BRANCED 1/CYCLOIDEA/PCF (TCP) 14 and TCP15 proteins (Peng et al., 2015) which repress endoreduplication by transcriptional control of *RETINOBLASTOMA-RELATED1* (*RBR1*) and *CYCLIN A2;3* (*CYCA2;3*) gene expression (Li et al., 2012).

Here we show that DA1 is an endopeptidase activated by multiple ubiquitylation mediated by the E3 ligases BB and DA2. In a feedback mechanism, DA1 then cleaves BB and DA2, leading to their destabilization. DA1-mediated cleavage of BB exposed a destabilizing N-terminal that was substrate for the N-end rule E3 ligase PROTEOLYSIS 1 (PRT1). This mechanism is predicted to transiently activate DA1 peptidase, which also cleaves of UBP15, TCP15 and the related TCP22, leading to their predicted inactivation and de-stabilization. DA1 peptidase may therefore contribute to the concerted transition from cell proliferation to endoreduplication and differentiation, limiting organ size.

## RESULTS

### Genetic and physical interactions of DA1, BB and DA2

We previously identified genetic interactions between the *da1-1* allele of *DA1* and genes encoding the RING E3 ligases BB (Li et al. 2008) and DA2 (Xia et al. 2013) that led to synergistic increases in seed and organ sizes. In this study we use the *da1-1* enhancing allele of *BB* called *eod1-2* (Li et al. 2008) and refer to the mutant version as *bb-eod1-2* and the wild-type version of as *BB*. The *da1-1* allele, an R358K change in a highly conserved region, had a negative influence on the functions of *DA1* and the close family member *DAR1*, but the basis of this was not known, which complicated interpretation of DA1 function. We therefore assessed phenotypes of a loss-of function T-DNA allele of *DA1* (*da1-ko1*).

Measurements of petal and seed sizes using high-resolution scanning showed that the *da1-ko1* T-DNA allele led to increased petal and seed sizes (Fig. 1A,B), and it also interacted genetically with the loss-of-function allele *bb-eod1-2* and *da2-1* in both petal size and seed area. This showed that *DA1* can be studied independently of other *DA1* family members. Both the *da1-1* and *bb-eod1-2* mutations increased the maximum growth rate, while the double mutant *da1-1 bb-eod1-2* showed a further increased maximum growth rate and continued to growth for approximately 5 days longer than either single mutant (Fig. 1C). The time at maximum growth rates was slightly earlier in *bb-eod1-2* than Col-0, in contrast to *da1-1* and *da1-1 bb-eod1-2*, which showed a 3-day retardation of the time of maximum growth rate, and final leaf sizes showed a more than additive increase in the double mutant, as observed previously (Li et al. 2008). These data indicated that *BB* may influence leaf final size at earlier stages of growth than *DA1*. We previously demonstrated that DA1 and DA2 physically interact (Xia et al. 2013). Pull-down experiments showed that GST-tagged DA1 also interacted with HIS-tagged BB, but not with HIS-tagged BBR (Big Brother Related, At3g19910), a close homolog of BB (Breuninger and Lenhard 2012) (Fig. 1D). These *in vitro* interactions were verified by *Agrobacterium*-mediated co-expression of BB-GFP and Myc-tagged DA1 in *Nicotiana benthamiana* leaves. Myc-DA1 was only detected in a complex with BB-GFP, and not GFP (Fig. 1E).

**Figure 1.**
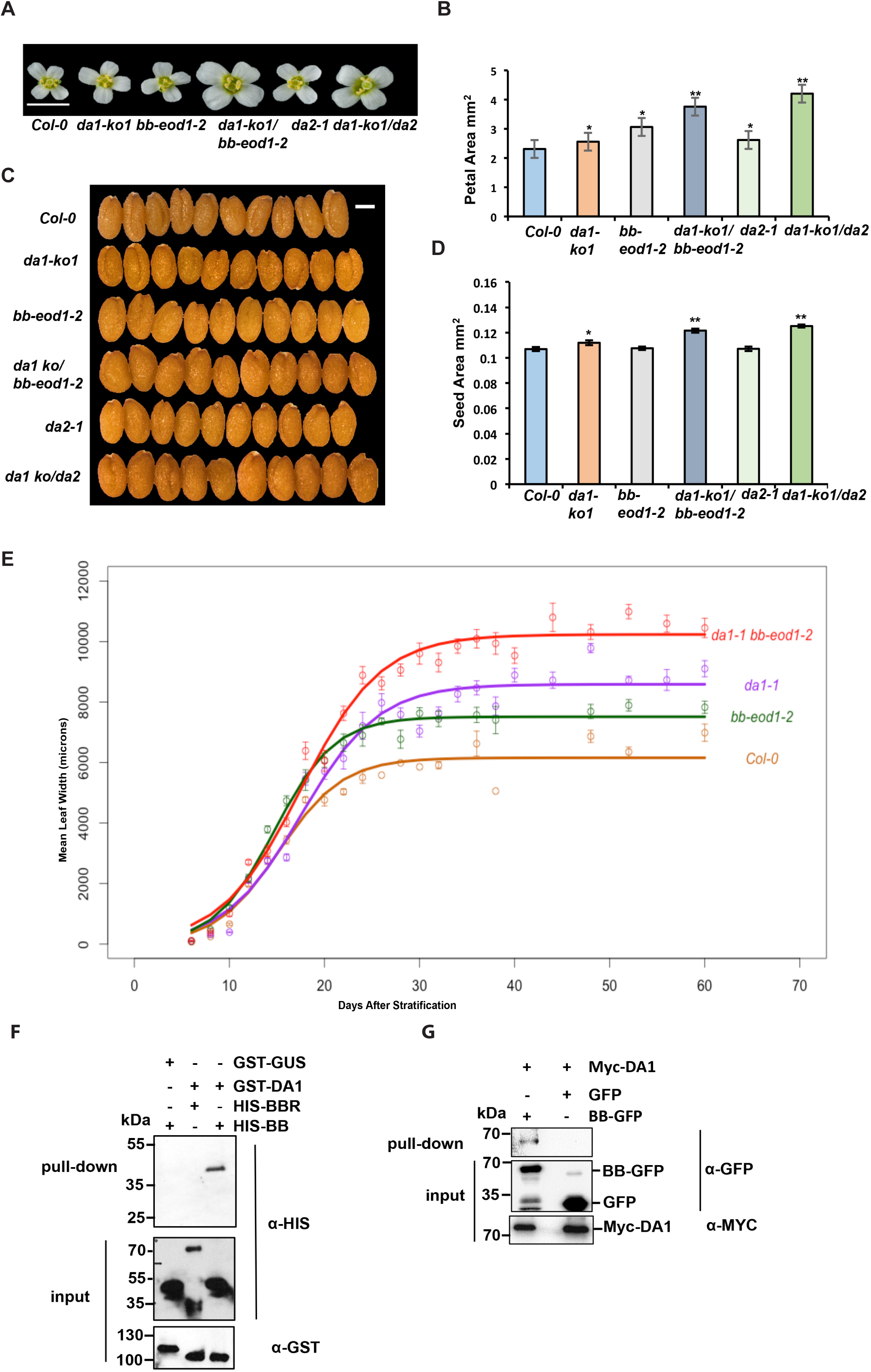
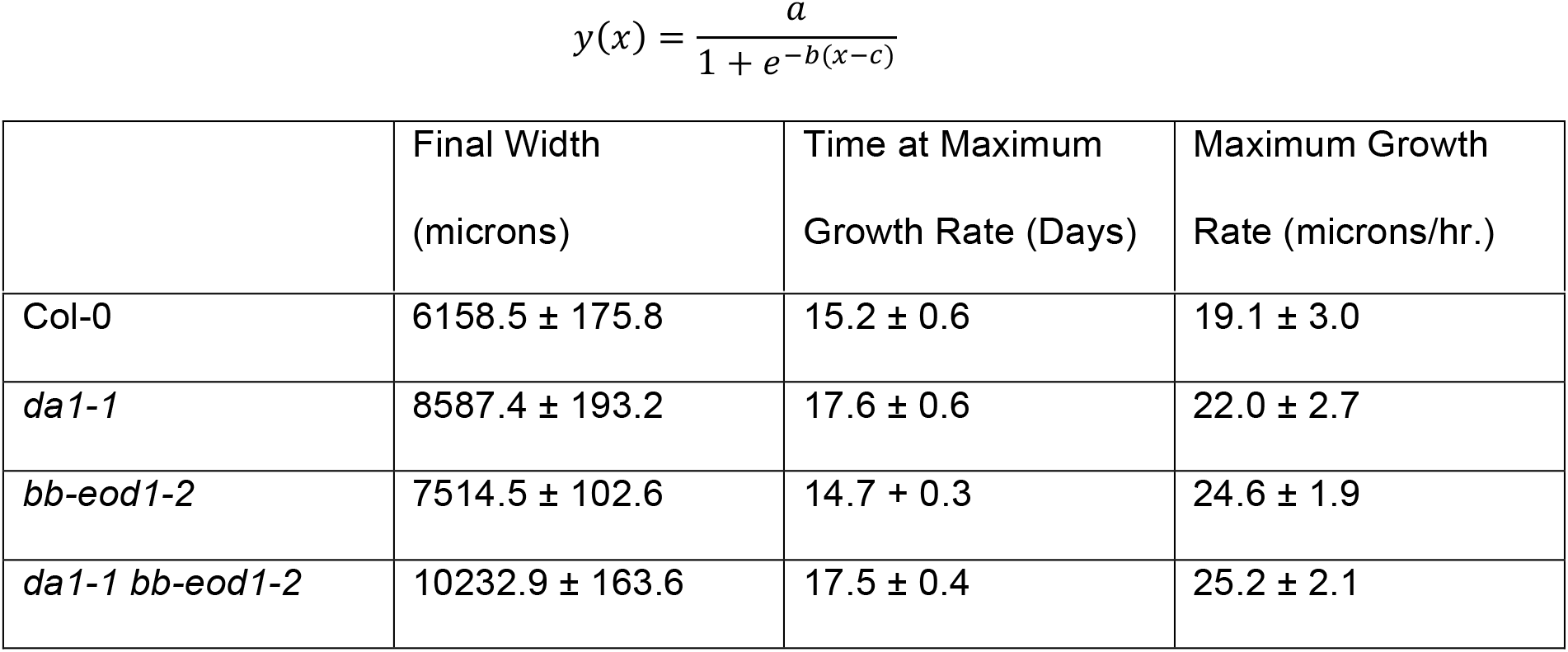
mGenetic and Physical Interactions of DA1, BB and DA2. (A and B) The single loss-of-function *da1-ko1* allele interacts with *bb-eod1-2* and *da2-1* to increase petal area. Panel A is an image of flower heads showing the sizes of petals. Scale bar is 5 mm. Panel B shows petal areas. Values given are means (n=36) ± SE. *P<0.05, **P<0.01 (Student’s *t* test) compared to wild type Columbia. (C and D) The single loss-of-function *da1-ko1* allele interacts with *bb*-*eod1-2* and *da2-1* to increase seed area. Panel C shows 10 seeds aligned to reveal size differences. Scale bar is 2mm. Panel D shows seed areas. Values give are means (n=50) ± SE. *P<0.05, **P<0.01 (Student’s *t* test) compared to wild type Columbia. (E) Dynamic growth measurements of leaf 1 width in Col, *da1-1* and *da1-1 bb-eod1-2*. Lines were fitted to data points using the sigmoidal function (see Supplemental Methods) (F) DA1 interacts with BB *in vitro*. GST-DA1 interacted with HIS-BB. GST-DA1 did not interact with HIS-BBR, an E3 ligase closely related to BB. GST-GUS (β-glucuronidase) was used a negative control. (G) Myc-Tagged DA1 interacted with BB-GFP after transient co-expression in *Nicotiana benthamiana* leaves. BB-GFP and GFP were co-expressed with Myc-DA1 using *Agrobacterium*-mediated transient expression in *N. benthamiana* leaves. Expressed proteins were purified using GFP-trap and immunoblotted.

### DA1 is multiply ubiquitylated by BB and DA2

The interactions of DA1 with BB and DA2 suggested that DA1 might be a substrate of these RING E3 ligases, so we conducted *in vitro* ubiquitylation reactions using BB, DA2 and BBR (BB-Related) E3 ligases. Fig. 2A shows that BB ubiquitylated DA1 in an E1- and E2-dependent reaction, as did DA2 (Fig. 2B), while BBR did not (Fig. 2C). The extent of DA1 ubiquitylation suggested that DA2 was more efficient at ubiquitylation than BB, and the sizes of ubiquitylated DA1 indicated that between 4-7 ubiquitin molecules may be conjugated to DA1. Mass spectrometric analyses of ubiquitylated DA1 prepared *in vitro* was used to identify peptides containing the characteristic di-glycine ubiquitylation signature of a lysine residue (KGG). Analysis of DA1 ubiquitylated by DA2 or BB identified seven ubiquitylated lysine residues in DA1, with 4 lysines in the C-terminal domain of DA1 (K381, K391, K475 and K591) consistently conjugated with ubiquitin (Supplemental Fig. S1). This number of ubiquitylation sites concurred with the patterns of ubiquitylation observed in Fig. 2A,B, suggesting that DA1 molecules are multiply ubiquitylated (Haglund et al. 2003; Komander and Rape 2012). Mutation of the consistently ubiquitylated lysines to arginine in DA1 (termed DA1(4K-4R)) did not reduce ubiquitylation by DA2 *in vitro* (Fig. 2D), and mass spectrometric analyses showed ectopic ubiquitylation of other lysines across DA1 (Supplemental Fig. S2). Therefore, the DA1 ubiquitylation mechanism has a preference, but not specificity, for certain lysines. These patterns of ubiquitylation are shown in Fig. 2D.

**Figure 2.**
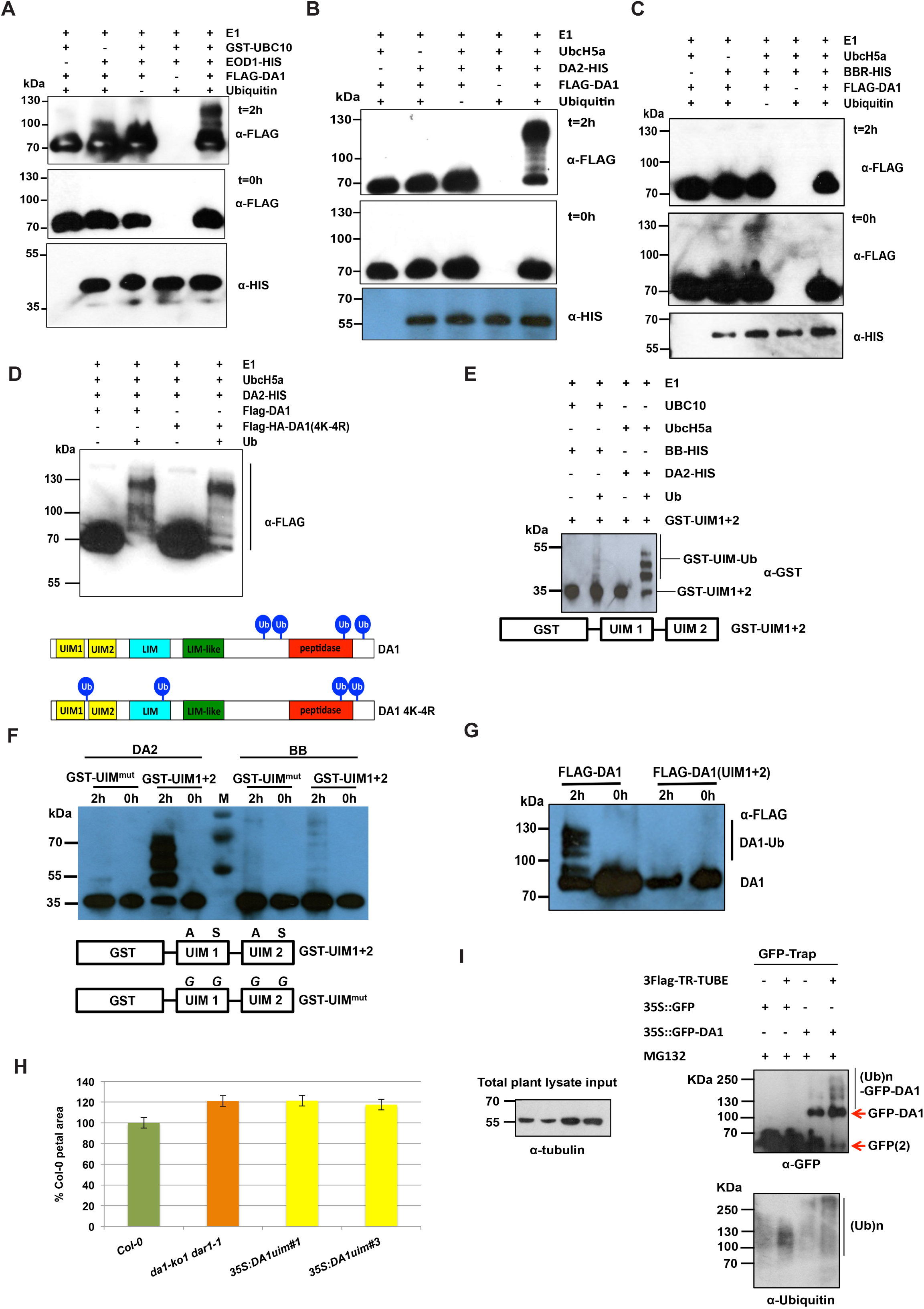
DA1 is multiply ubiquitylated by BB and DA2 in a UIM-dependent reaction. (A, B and C) *in vitro* ubiquitylation of DA1 by the RING E3 ligases BB (panel A) and DA2 (panel B), but not by BBR (panel C) in an E1-, E2- and ubiquitin-dependent reaction. Anti-FLAG antibodies detected FLAG-ubiquitylated forms of FLAG-DA1 ranging from above 70 kDa to approximately 130 kDa. Anti-HIS antibodies detected BB-HIS, DA2-HIS or BBR-HIS fusion proteins. (D) FLAG-DA1 and FLAG-DA1(4K-4R) are both ubiquitylated by DA2 in similar patterns in an *in vitro* ubiquitylation reaction. In the lower panel Ub represents a ubiquitin moiety conjugated to a lysine at the approximate location in DA1 and DA1(4K-4R). Regions of protein similarity with known domains are shown: UIM1 and UIM2 are similar to Ubiquitin Interaction Motifs; LIM is similar to canonical LIM domains, LIM-like is a related motif found in DA1 family members, and peptidase contains a predicted peptidase active site. (E) An *in vitro* ubiquitylation reaction with DA2 and BB as E3 ligases and GSTUIM1+2. GST-UIM1+2 is ubiquitylated in a similar pattern to DA1 by both DA2 and BB, with DA2 conferring higher levels of ubiquitylation than BB. (F) An *in vitro* ubiquitylation reaction with DA2 and BB as E3 ligases and GST-UIM1+2 with mutations that reduce ubiquitin binding. Mutated versions of UIM1 and UIM2 strongly reduced DA2- and BB-mediated ubiquitylation of GST-UIM1+2. (G) A time-course of FLAG-DA1 and FLAG-DA1(UIM1+2) with mutations in the UIMs as in panel E. These strongly reduced DA1 ubiquitylation. (H) DA1(UIM1+2) is not functional *in vivo* as it does not complement the large petal phenotype of the *da1-ko1 dar1-1* double mutant. Two independent homozygous T-DNA insertion lines were scored for petal size and compared to wt Col-0 and *da1-ko1 dar1-1*. Values given are means (n=120) ± SE, expressed as percentage of wild-type Col-0 petal areas. Student’s *t* test showed no significant differences between the transformants and the parental *da1-ko1 dar1-1* line. (I) Transgenic Arabidopsis plants expressing a GFP-DA1 fusion protein under control of the 35S promoter were used to detect DA1 ubiquitylation *in vivo*. GFP runs as a dimer on the gel due to high protein concentrations. Protein extract input levels are shown using anti-tubulin antibody.

DA1 and four other family members have multiple UIMs that interact with ubiquitin (Li et al. 2008; Peng et al. 2015). UIMs are part of a larger class of Ubiquitin Binding Domains (UBD) formed from a single α helix that is often found in multiple arrays (Hicke et al. 2005; Husnjak and Dikic 2012). Tandem UIMs have been shown to bind K63-linked ubiquitin chains in the mammalian DNA repair protein RAP80 (Sato et al. 2009). To assess their function in DA1, the N-terminal region of DA1 containing mutated UIM1 and UIM2 was fused to GST and expressed in *E. coli*, and conserved Ala and Ser residues, predicted to be in the α helical domain of the UIMs (Kim et al. 2007) (Supplemental Fig. S2), were mutated to Gly in both UIMs of GST-UIM1+2 and in DA1. GST-UIM1+2 bound ubiquitin and mutation of UIM1 alone did not reduce binding of ubiquitin, while mutation of UIM2 abolished ubiquitin binding, confirming that the GST-UIM1+2 protein bound ubiquitin *via* its UIM motifs (Supplemental Fig. S2). Fig. 2E showed UIM1+2 conferred BB- and DA2-dependent ubiquitylation on GST *in vitro*, with DA2 again facilitating higher levels of ubiquitylation. Fig. 2F showed that mutation of both UIM1 and UIM2 in GST-UIM1+2 strongly reduced *in vitro* ubiquitylation of GST-UIM1+2 by BB and DA2. The UIM1 and UIM2 mutations in DA1 also reduced its ubiquitylation *in vitro* (Fig. 2G), and DA1 with mutated UIMs did not complement the large petal size in the double mutant *da1-ko dar1-1* by (Fig. 2H). To detect ubiquitylation *in vivo*, DA1 was expressed from the constitutive 35S promoter as an N-terminal GFP fusion protein and purified from seedling tissues using a GFP-Trap. Characteristic patterns of DA1 ubiquitylation were detected on purified GFP-DA1 (Fig. 2I, right panel). Therefore DA1 is ubiquitylated by the E3 ligases BB and DA2 *in vitro* by a UIM1 and UIM2 – dependent mechanism, DA1 is ubiquitylated *in vivo*, and UIMs were required for DA1 function.

### DA1 cleaves BB and DA2 with a ubiquitin-dependent peptidase activity

A time-course of BB-HIS incubated with purified FLAG-DA1 that had been ubiquitylated by BB, or incubated with non-ubiquitylated FLAG-DA1, showed that in the presence of ubiquitylated DA1 a HIS-tagged BB fragment of approximately 35 kDa was produced after 4 h incubation (arrowed in Figure 3A). When ubiquitylated FLAG-DA1 was incubated with DA2-HIS, a 25 kDa HIS-tagged DA2 cleavage product was also detected after 4 h incubation (arrowed in Fig. 3A). Similar experiments using FLAG-DA1 ubiquitylated by DA2 showed identical patterns of BB-HIS and DA2-HIS cleavage (Fig. 3B). BBR-HIS did not show a cleavage product in these conditions. Thus DA1, ubiquitylated by either BB or DA2, generated cleavage products from both BB and DA2 *in vitro*.

**Figure 3.**
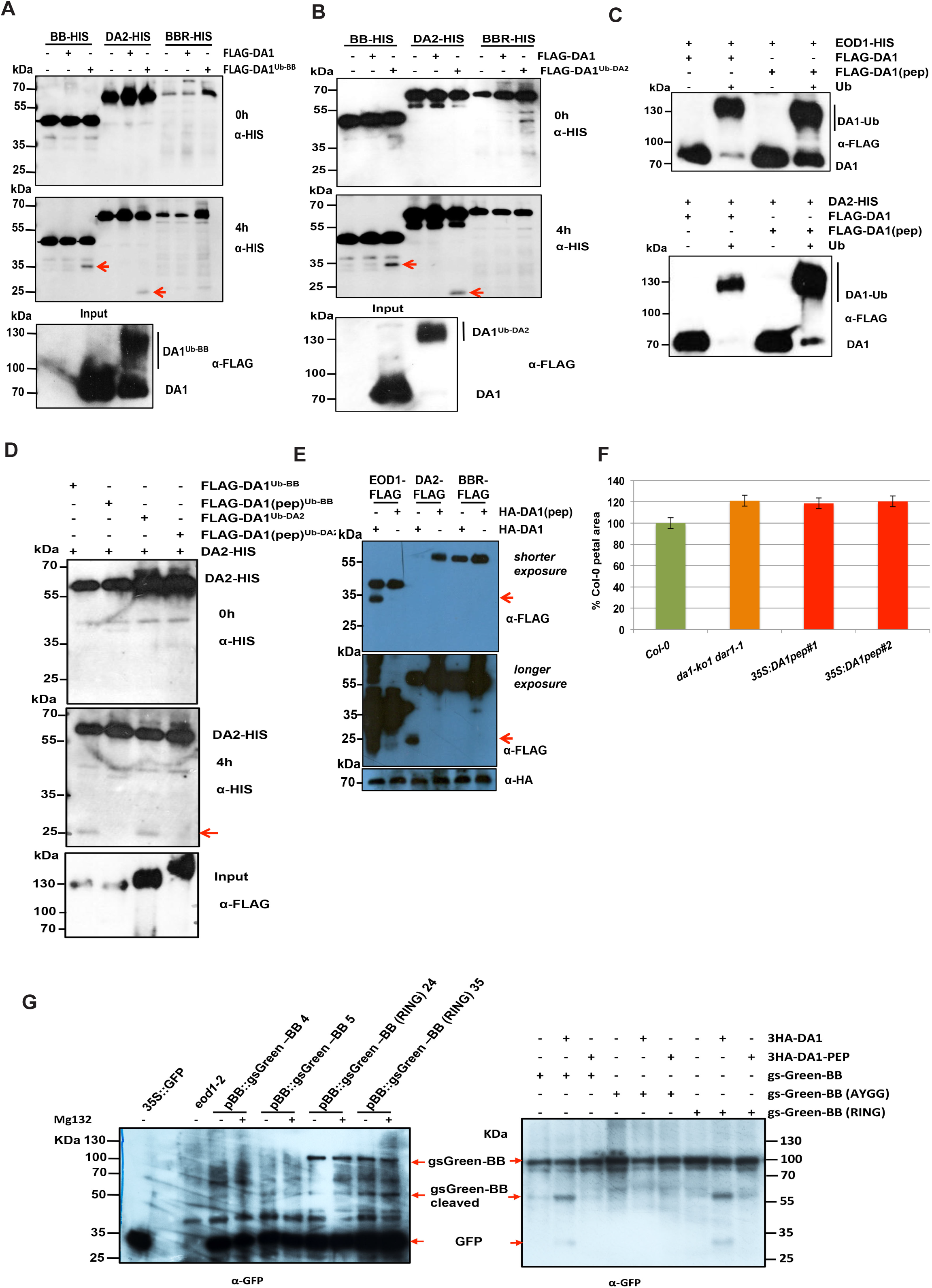
DA1 is an endopeptidase activated by multiple ubiquitylation, and cleaves the E3 ligases BB and DA2 that ubiquitylate it. (A and B) Time-course of an *in vitro* reaction of FLAG-DA1 or FLAG-DA1 ubiquitylated by BB (panel A) or DA2 (panel B) with BB-HIS, DA2-HIS and BBR-HIS. The lower panels show loading of FLAG-DA1 and FLAG-DA1^Ub-BB^. After 4h, cleavage products (shown by red arrows) of BB and DA2 had been produced by ubiquitylated FLAG-DA1, but not by FLAG-DA1. BBR was not cleaved under these conditions. (C) An *in vitro* ubiquitylation reaction of DA1 and DA1(pep) using BB-HIS (upper panel) and DA2-HIS (lower panel) as E3 ligases. Ubiquitin-dependent multiple monoubiquitylation of FLAG-DA1 and FLAG-DA1(pep) by both BB and DA2 was detected. (D) A time-course of an *in vitro* cleavage reaction using DA2-HIS as a substrate (left panels), and FLAG-DA1 or FLAG-DA1(pep) ubiquitylated by either BB or DA2 (loading shown on the bottom panel). The red arrow in the lower left panel indicates the DA2 cleavage product at 4h that was only produced by FLAG-DA1^Ub^ and not by FLAG-DA1(pep)^Ub^. (E) Arabidopsis *da1-ko1 dar1-1* mesophyll protoplasts were co-transfected with plasmids expressing BB-FLAG, DA2-FLAG, BBR-FLAG, and HA-DA1 and HA-DA1(pep). The same sized cleavage products (red arrows) from BB-FLAG and DA2-FLAG were detected as seen in panels (A) and (B) above. Longer exposure of the top immunoblot (central panel) showed cleaved DA2. The lower panel shows loading of HA-DA1 and HA-DA1(pep). (F) DA1(pep) is not functional *in vivo* as it does not complement the large petal phenotype of the *da1-ko1 dar1-1* double mutant. Transformants expressing *35S::DA1(pep)* were scored for petal size and compared to wt Col-0 and *da1-ko1 dar1-1*. Values give are means (n=150) ± SE, expressed as percentage of wild-type Col-0 petal areas. Student’s *t* test showed no significant differences between the transformants and the parental *da1-kodar1-1* line. (G) Cleavage of gsGreen-BB is shown *in planta* (left panel) and in transiently expressed protoplasts in the right panel for comparison. Large scale protein extracts from transgenic eight day old seedlings expressing *BB::gsGreen-BB* and *BB::gsGreen*-BB (RING) were purified on a GFP-trap. Loading controls used levels of free GFP. The expected size cleavage product (arrowed) was observed in plant extracts and in protoplasts for comparison.

Examination of the conserved C-terminal region of DA1 revealed an extended sequence motif, HEMMHX_15_EE (Supplemental Fig. S3), which is a zinc aminopeptidase active site found in clan MA endopeptidases (Rawlings et al. 2012). The HEMMH motif was mutated to AEMMA, removing the putative Zn-coordinating histidine residues, to form DA1(pep). Fig. 3C showed that DA1(pep) and DA1 were ubiquitylated *in vitro* to an equal extent by both BB and DA2. In an *in vitro* time-course reaction, ubiquitylated DA1(pep) did not generate the 25 kDa HIS-tagged DA2 band seen after incubation with ubiquitylated DA1 (Fig. 3D). Co-expression of BB-FLAG, DA2-FLAG or BBR-FLAG with HA-DA1 or HA-DA1(pep) in *da1-ko1 dar1-1* mutant leaf protoplasts showed that HA-DA1 but not HA-DA1(pep) generated a similar-sized 35 kDa BB-FLAG cleavage product (arrowed in the top panel of Fig. 3E) as seen in *in vitro* reactions (Fig. 3A,B). Longer exposure of the same Western blot (bottom panel of Figure 3E) was required to identify the 25 kDa DA2-FLAG cleavage product, which was not generated by co-expression with DA1(pep). Fig. 3F shows the mutation in *DA1* abolishing DA1 peptidase activity did not complement the *da1-ko1 dar1-1* large petal phenotype, establishing that DA1 peptidase activity is required for *in vivo* function. To detect DA1 peptidase activity *in vivo*, transgenic plants expressing a BB::gsGreen-BB fusion protein, and a RING-domain mutant version that was predicted to be more stable *in vivo* due to reduced auto-polyubiquitylation (Disch et al. 2006a) were generated. Analysis of GFP-Trap purified proteins (Fig. 3G, left panel) showed a cleavage product of the expected size generated from RING-mutant gsGreen protein in two independent transformants. Full-length wild-type gsGreen-BB was not detected, although low levels of an expected cleavage product were identified. For comparison, the same constructs, together with a non-cleavable form (AY-GG, see Fig. 5B) were expressed using the 35S promoter in protoplasts with DA1 (Fig. 3G, right panel). This showed the predicted DA1-mediated BB cleavage product, which was not generated in the AY-GG version of BB.

A Förster Resonance Energy Transfer (FRET) DA1 peptidase sensor was constructed using eGFP donor and mCherry acceptor pairs (van der Krogt et al. 2008) connected by BB to provide another measure of DA1 peptidase activity *in vivo*. Cleavage of the fluorophore pair by DA1 would increase the fluorescence lifetime towards that of eGFP-BB compared to that of the intact sensor protein by impairing energy transfer between the fluorophores. The peptidase sensor and a control donor sensor were transfected into *da1-ko1 dar1-1* root protoplasts and Fluorescence Lifetime Imaging (FLIM) was performed. Fig. 4A shows that the fluorescence lifetime (τ) of the GFP-BB donor control was approximately 2.48 ns, while that of an intact donor-acceptor pair was approximately 2.25 ns, demonstrating efficient FRET. When co-transfected with DA1, the fluorescent lifetime of the donor-acceptor pair increased to approximately 2.38 ns. Lifetime imaging of typical transfected protoplasts showed a generalized cellular localization of DA1-mediated cleavage. Fig. 4B showed the eGFP-BB-mCherry donor-acceptor pair was cleaved by DA1 peptidase at the expected site in transfected root protoplasts. Therefore, DA1 has a latent peptidase activity that is activated by multiple ubiquitylation mediated by its UIM 1+2 domain and the RING E3 ligases BB and DA2, and activated DA1 peptidase then specifically cleaves these two E3 ligases.

**Figure 4.**
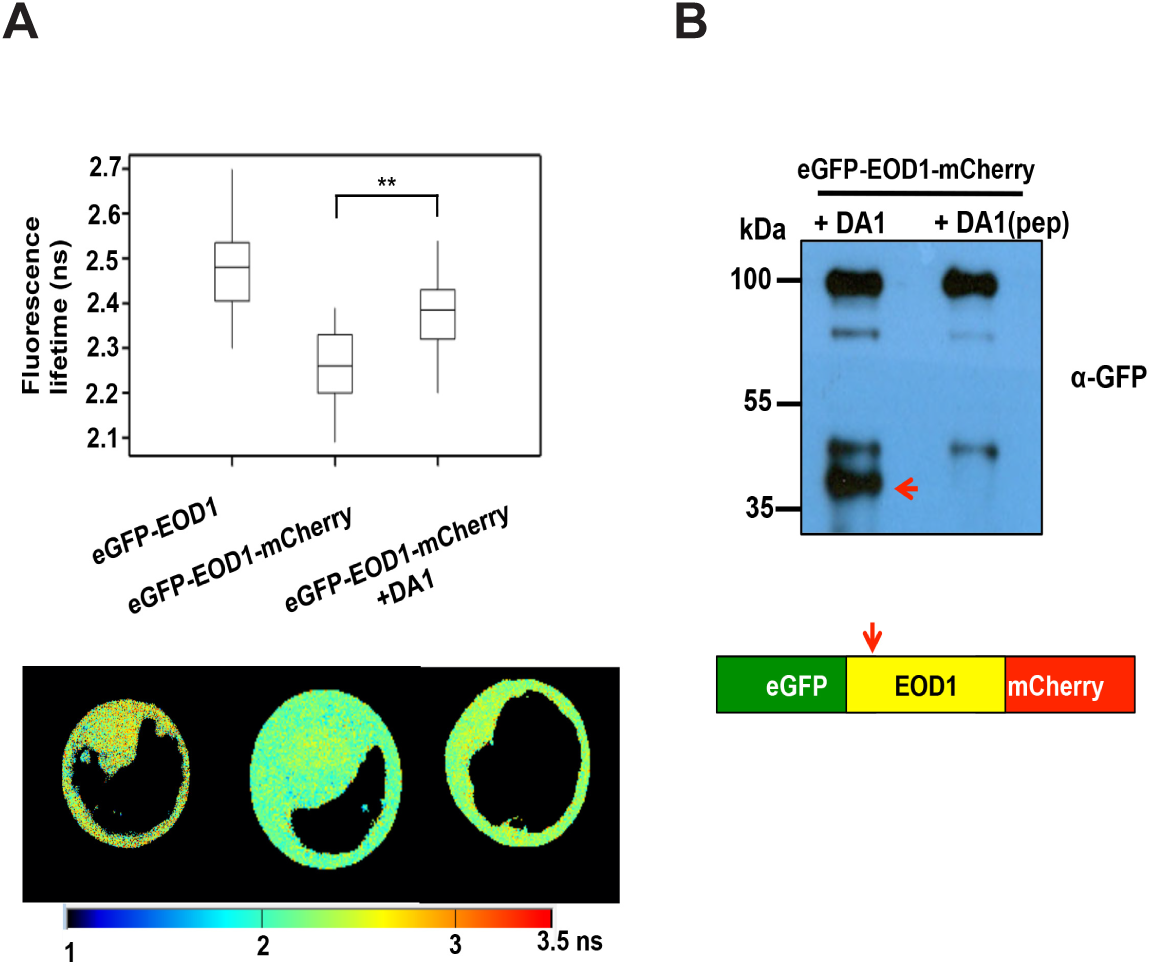
Detection of DA1-mediated cleavage of BB *in vivo* using FRET. (A) Root protoplasts of *da1-ko1 dar1-1* plants were transfected with the FRET construct eGFP-BB, or a control eGFP-BB-mCherry construct, together with DA1 to detect DA1-mediated cleavage of BB. Transfected protoplasts were imaged using multi-photon microscopy and fluorescence half-times of protoplasts (n= 13) were captured. The heat-map shows fluorescent lifetime values and typical protoplasts are shown to illustrate fluorescent half-lives imaged over the cell. The box plots show significantly increased fluorescence lifetime after DA1 transfection (**; *P* ≤ 0.001, Student’s *t* test). (B) Cleavage of eGFP-BB-mCherry by DA1 in the imaged protoplasts shown in (A). The arrow shows the major cleavage product of approximately 40 kDa expected from DA1 cleavage near the N-terminus of BB.

### Identification of a DA1 peptidase cleavage site in BB

To define the potential functions of DA1-mediated cleavage, the DA1 cleavage site in BB was identified using Edman sequencing of purified cleaved BB-HIS. Supplemental Fig. S4 shows neo-N terminal amino acid sequences that had a unique match to six amino acids in BB (Fig. 5A). This indicated a potential DA1 cleavage site within BB between A_60_ and Y_61_, consistent with the sizes of BB and its ca. 35 kDa cleaved form (Fig. 3A). Two mutant forms of BB were made to assess this potential DA1 cleavage site: a four amino acid deletion surrounding the site (ΔNAYK); and changing AY to GG (AY-GG) (Fig. 5B). These proteins were co-expressed in Arabidopsis *da1-ko1 dar1-1* mesophyll protoplasts as C-terminal FLAG fusion proteins with HA-DA1 and HA-DA1(pep). Fig. 5B showed that the mutant BB-FLAG proteins were not cleaved by DA1, establishing that DA1 peptidase activity cleaved BB between A_60_ and Y_61_. A cleaved form of BB called MY61-BB was also made with an initiator Met followed by Y_61_ (Fig. 5B). MY61-BB was expressed using the 35S promoter in *da1-ko1 bb-eod1-2* mutant Arabidopsis. Its lack of complementation of *bb-eod1-2* (Fig. 5C) showed that DA1 peptidase-mediated cleavage reduced BB activity.

**Figure 5.**
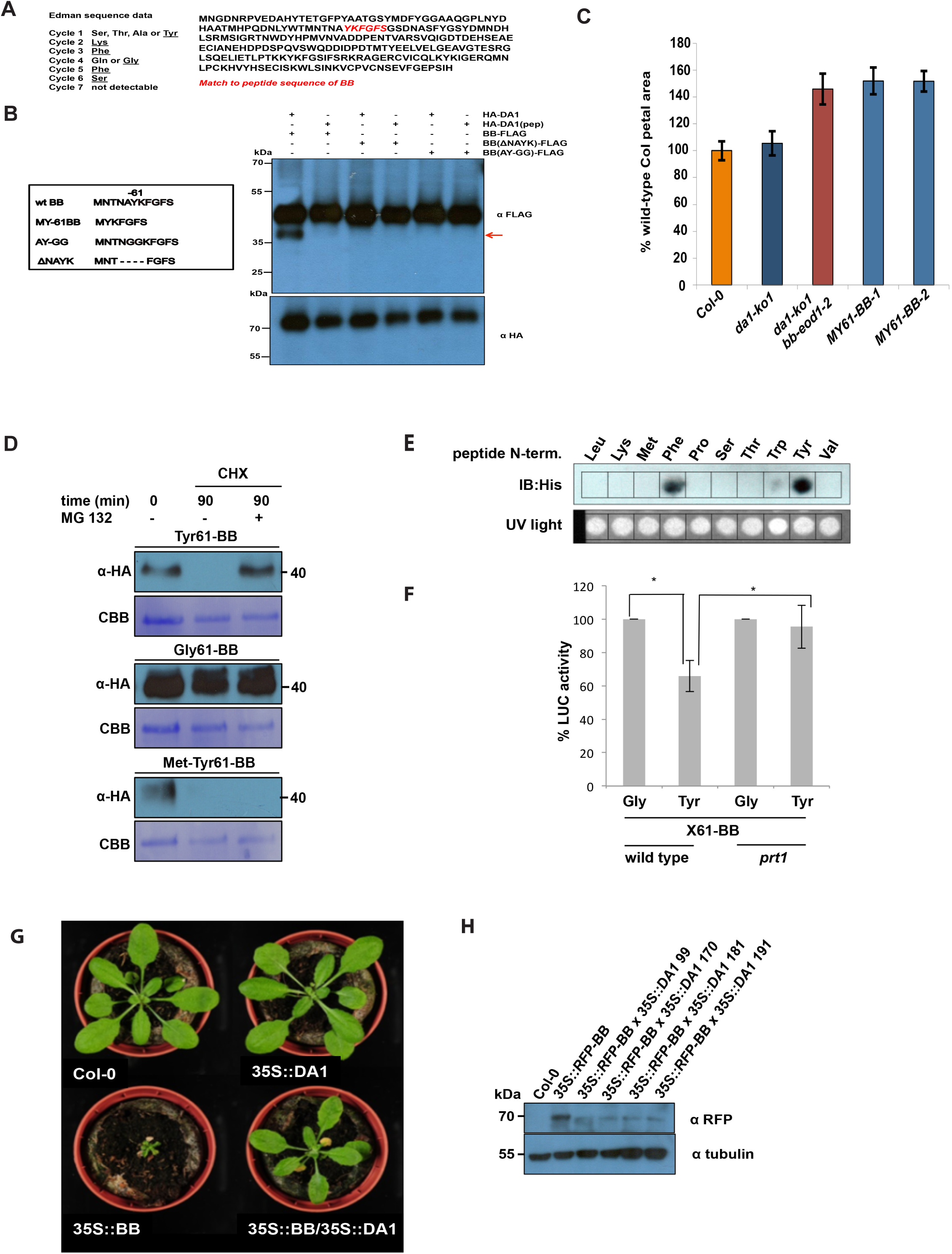
Identification of the DA1 cleavage site in BB and destabilization and functional inactivation of cleaved BB *in vivo* by the N-end rule. (A) Neo-N terminal sequences of purified cleaved BB-HIS (left) matched the complete BB protein sequence (right, shown in red). Data from six Edman sequencing cycles is shown in Figure S5. (B) The predicted DA1 cleavage site in BB was mutated by changing the amino acids flanking the site AY to GG (AY-GG), and by deleting 2 amino acids from both sides of the predicted cleavage site (ΔNAYK). BB-FLAG, BB (ΔNAYK)-FLAG, and BB (AY-GG)-FLAG were expressed in *da1-ko1 dar1-1* Arabidopsis mesophyll protoplasts under control of the by the 35S promoter. HA-DA1 did not cleave BB containing mutations in the predicted cleavage site. The lower panel shows HA-DA1 and HA-DA1(pep) loading. (C) A DA1-cleaved version of BB does not function *in planta*. A cleaved version of BB termed MY61-BB was expressed in *da1-ko1 bb-eod1-2* plants under control of the 35S promoter. The graph compares petal areas in wt Col, *da1-ko1*, *da1-ko1 bbeod1-2* and two independent transgenic lines. Values give are means (n=50) ± SE, expressed as percentage of wild-type Col petal areas. Student’s *t* test showed no significant differences between the transformants and the parental *da1-ko1 bb-eod1-2* line. (D) *In vitro* degradation of BB is dependent on N termini. Ubiquitin-fusion constructs were expressed in a reticulocyte lysate cell-free system. Samples were incubated for 30 mins with or without MG132. Then cycloheximide (CHX) was added to inhibit translation and samples were taken 0 and 90 mins after CHX addition. Samples were electrophoresed on SDS-PAGE and immunoblotted used HA antibodies to detect BB protein levels. Loading controls are the CBB stained membrane. (E) Constructs expressing ubiquitin fusions of -61BB-Luciferase (Ub-Gly-61BB-HA-LUC and Ub-Tyr-61BB-HA-LUC), co-translationally generating glycine or tyrosine neo-N-termini, were transformed into Col-0 or *prt1* mutant protoplasts. Transformation efficiency was measured using *pUBC::GUS* control plasmid. Luciferase activities were normalized to GUS activity, and activity of Gly-61BB-HA-LUC was taken as 100%. Three independent transformation experiments were conducted. The significance of differences was calculated using Student’s *t*-tests (two sites, uncoupled). * p-value ≤ 0.05. (F) Constructs expressing ubiquitin fusions of -61BB-Luciferase constructs with glycine or tyrosine neo N-termini (Ub-Gly-61-BB-HA-LUC and Ub-Tyr-61-BB-HA-LUC) were transfected into wt or *prt1* mutant protoplasts. Transfection efficiency was measured using a pUBC::GUS control. Luciferase activities were normalized to GUS activity, and the LUC activity of Gly-61-BB-HA-LUC was taken as 100%. From three independent transformation experiments, the significance of differences was calculated using Student’s *t*-tests (two sites, uncoupled). * p-value ≤ 0.05. (G) Over-expression of BB under control of the 35S promoter leads to strongly reduced growth, and when crossed with a line over-expressing DA1, this growth inhibition was reversed. This demonstrated that DA1 can reduce the growth inhibitory effect of high levels of BB. (I) Crossing a line over-expressing DA1 into a line expressing 35S::RFP-BB reduced RFP-BB levels. Homozygous progeny of four independent crosses (99, 170, 181 and 191) are shown.

### BB stability is dependent on it N-terminus and N-end rule function

DA1 cleavage products of DA2 were unstable, indicating that one function of DA1-mediated cleavage may be to destabilize proteins (Fig. 3E). This was also observed for BB in cell-free degradation assays, in which MY61-BB was unstable compared to wild-type BB (Supplemental Fig. S5). To test the role of the neo-N terminus of BB on protein stability, -61BB proteins with different N-termini (Y, G, MY) were expressed using the ubiquitin fusion technique (UFT) (Bachmair et al. 1986). HA-tagged constructs were translationally co-expressed in a cell-free rabbit reticulocyte system, with or without MG132 proteasome inhibitor, and translation stopped by the addition of cycloheximide. Y61-BB was highly unstable, whereas G61-BB was stable (Fig. 5D). Interestingly, the artificial MY61-BB was also highly unstable in a proteasome-independent mechanism. The neo-N terminal sequence of DA1-cleaved BB starts with YK, a potentially destabilizing sequence of a type II N-end rule degron (Varshavsky 2011). The N-end rule E3 ligase PROTEOLYSIS 1 (PRT1) mediates stability of model N-end rule substrates with such aromatic amino-terminal residues (Potuschak et al. 1998). To assess the potential role of PRT1 in N-end rule mediated degradation of BB, we tested the binding of PRT1 to 17-mer peptides representing variants of the neo-N termini of BB on a backbone sequence of an N-end rule test substrate in SPOT assays (Synthetic Peptide arrays On membrane support Technique). Purified recombinant HIS-MBP:PRT1 protein was incubated with the SPOT array and binding visualized by Western blotting. Recombinant PRT1 had a preference for binding to the large aromatic acids tyrosine and phenylalanine, consistent with previously suggested specificity (Fig. 5E) (Potuschak et al. 1998; Stary 2003; Faden et al. 2016). To assess whether PRT1 had a role in DA1-mediated BB degradation, BB was expressed with an N-terminal ubiquitin fusion and a C-terminal luciferase fusion to reveal neo-N termini in Col-0 or *prt1* mutant mesophyll protoplasts. BB-luciferase (LUC) activity was reduced in wild-type protoplasts with a neo-N terminal tyrosine, which was not seen in *prt1* mutant protoplasts (Fig. 5F). Neo-N terminal glycine BB-LUC levels were not altered in either Col-0 or *prt1* mutant protoplasts. This indicated a strong dependence of Tyr-61BB stability on PRT1 activity. *In planta* evidence supporting the role of DA1 in reducing the growth inhibitory role of BB via N-end rule mediated degradation was shown by the suppression of growth reduction in a transgenic line over-expressing –*RFP-BB* from the 35S promoter by over-expression of DA1 (Fig. 5G). Western blots (Fig. 5H) confirmed that over-expression of *DA1* from the 35S promoter reduced *RFP-BB* protein levels.

### Functional analyses of DA1

We previously showed that the *da1-1* allele of *DA1* has a negative interfering phenotype with respect to the closely related family member *DAR1* (Li et al. 2008). The peptidase activity of the protein encoded by the *da1-1* allele, called DA1(R358K), which has an arginine to a lysine residue altered in a highly conserved C terminal region (Supplemental Fig. S4) was assessed. This mutation did not influence ubiquitylation of FLAG-DA1(R358K) (Fig. 6A) and neither did it create a site for ectopic ubiquitylation of FLAG-DA1(R358K) as determined by mass spectrometric analysis (Supplemental Fig. S1C). The peptidase activity of ubiquitylated FLAG-DA1(R358K) was qualitatively assessed *in vitro* and *in vivo* (using HA-DA1(R358K)) by comparison to wild-type DA1 peptidase activity (Fig. 6A,B). Both assays showed that DA1(R358K) had lower peptidase activity compared to DA1, suggesting regions of the conserved C-terminal region are required for peptidase activity and that the *da1-1* phenotype may be due to reduced peptidase activity. Fig. 6B also shows that DA1(4K-4R) which is ubiquitylated (Fig. 2E), had peptidase activity towards BB. This suggested that precise patterns of ubiquitylation are not required for activating DA1 latent peptidase activity.

**Figure 6.**
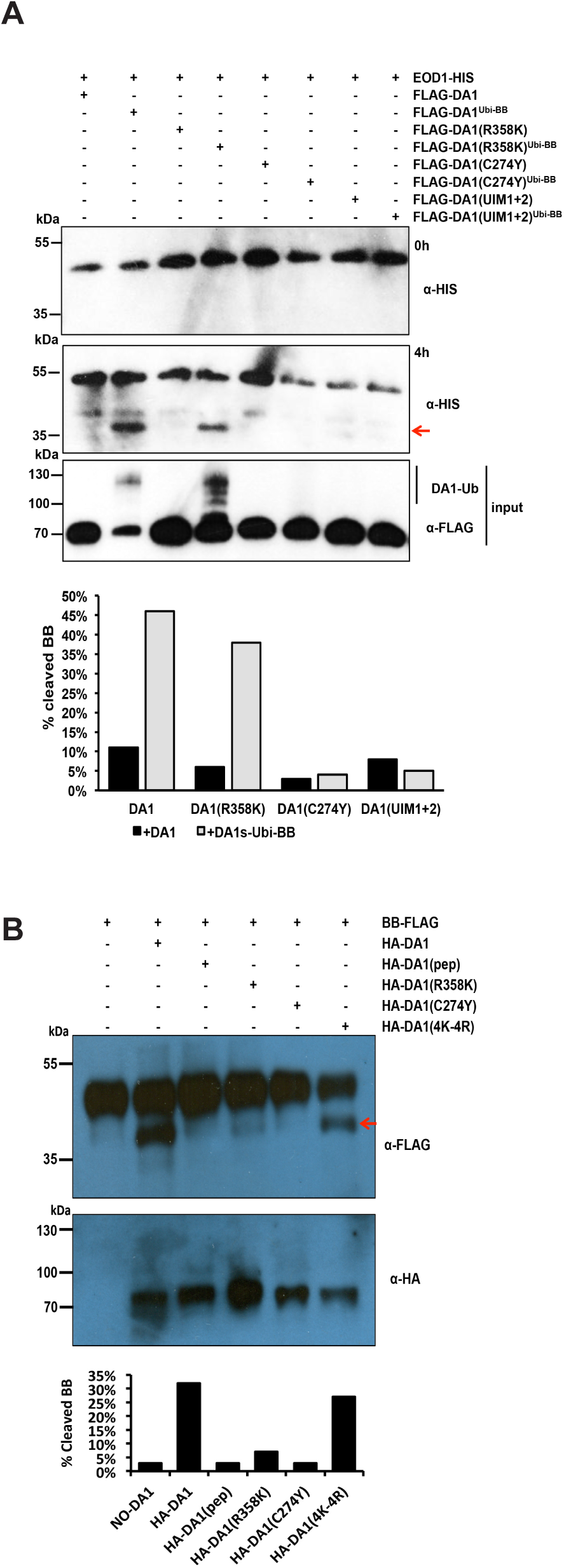
Functional analyses of DA1 activity *in vitro* and *in vivo*. (A) *In vitro* DA1 cleavage assays using BB-HIS as a substrate for ubiquitylated FLAG-DA1 and mutant versions. The top panel shows the reaction at 0h, and the middle panel shows the reactions at 4h. The red arrow indicates cleaved BB-HIS. The loading and ubiquitylation status of input HA-DA1 and mutant versions is shown. Quantitative scans showed BB-HIS was cleaved to a greater extent by wild-type DA1-HIS than by HA-DA1(R358K). HA-DA1(C274Y) and HA-DA1(UIM1+2) did not cleave BB-HIS. (B) *In vivo* DA1 cleavage reactions using BB-FLAG and HA-tagged DA1 and mutant versions. BB-FLAG cleavage product is arrowed. Quantitative scans showed reduced levels of BB cleavage by HA-DA1(R358K), and HA-DA1(C274Y) did not cleave BB-FLAG. HA-DA1(4K-4R) had normal levels of peptidase activity. The loading of HA-DA and its mutant versions is shown in the lower immunoblot.

*DAR4*, another *DA1* family member (Li et al. 2008), encodes a protein with an N-terminal TIR-NB-LRR, has a gain-of-function *chs3-2d* allele in the conserved C-terminal region (Supplemental Fig. S3) that activated constitutive defense responses (Xu et al. 2015). Alignments revealed high similarity to predicted protein sequences from the photosynthetic bacteria *Roseiflexus* sp (Burroughs et al. 2011) (Supplemental Fig. S6) that included four pairs of CxxC/H motifs with the potential to bind Zn similar to those in canonical LIM domains (Kadrmas and Beckerle 2004). The *chs3-2d* mutation changes a Cys to a Tyr in the third pair of conserved CxxC/H motifs (Supplemental Fig. S3 and S6), suggesting it may alter a possible LIM-like structure. This mutation was introduced into DA1 to create DA1(C274Y) and its activities assessed. Fig. 6A shows that DA1(C274Y) was not ubiquitylated by BB, and had no peptidase activity towards BB *in vitro* and *in vivo* (Fig. 6B). This implicated the putative LIM-like domain in DA1 in UIM-mediated ubiquitylation and activation of DA1 peptidase activity.

### DA1 peptidase activity cleaves TCP15, TCP22 and UBP15

The increased levels of UBP15 (Du et al. 2014), and TCP14 and TCP15 proteins (Peng et al. 2015) observed in the *da1-1* mutant suggested that DA1 activity may reduce the stability of these proteins by peptidase-mediated cleavage. Fig. 7A showed that DA1 peptidase cleaved UBP15 close to its C-terminus when transiently expressed together in *da1-ko1 dar1-1* protoplasts. The reduced signal in the western blot with the C-terminal FLAG fusion was due to the short FLAG-tagged protein running off the gel. TCP15 and the closely related TCP22 proteins were cleaved by DA1 in protoplasts (Fig. 7B), but we could not consistently detect TCP14 cleavage by DA1 or DAR1. These data show that DA1 peptidase activity can cleave UBP15, which promotes cell proliferation, and TCP15, which inhibits endoreduplication (Du et al. 2014; Peng et al. 2015).

**Figure 7.**
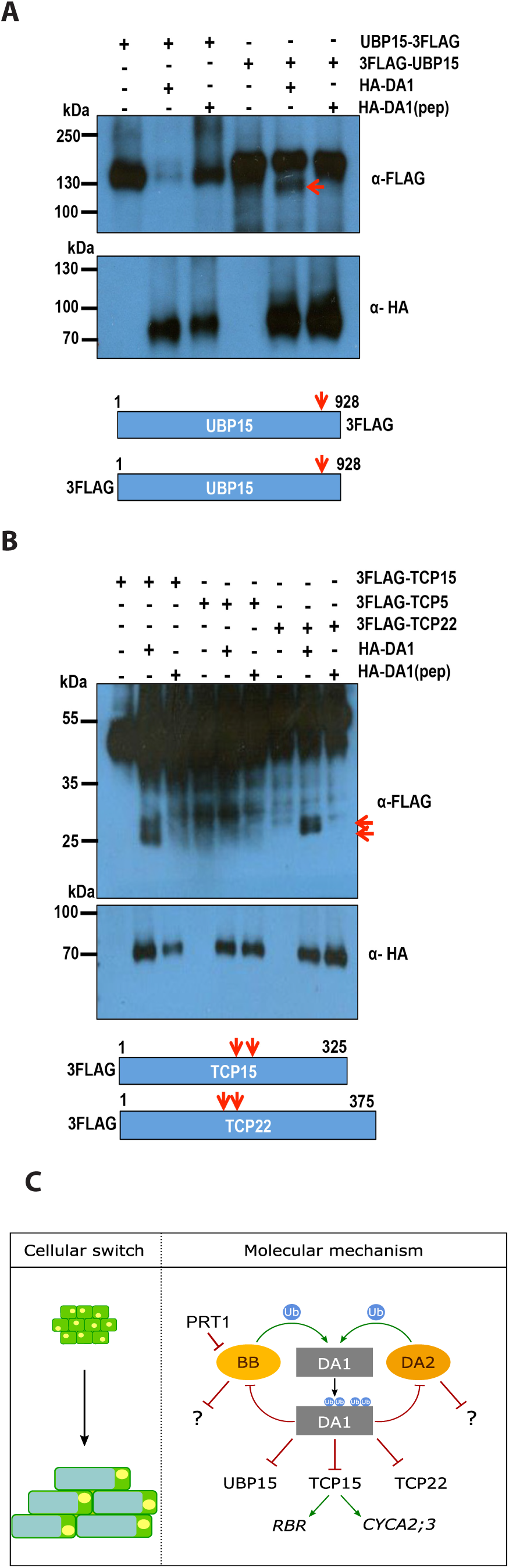
DA1 cleaves UBP15, TCP15 and TCP22 *in vivo*. (A and B) *In vivo* cleavage reactions of UBP15-3FLAG and 3FLAG-UBP15 (panel A), and 3-FLAG-TCP15, 3-FLAG-TCP22, and 3-FLAG TCP5 (a non-cleaved control) (panel B) using HA-DA1 and HA-DA1(pep). Constructs expressed from the 35S promoter were co-transfected into *da1-ko1 dar1-1* mesophyll protoplasts. UBP15, TCP14, TCP15, and TCP22 cleavage products are shown in the upper immunoblots. The lower immunoblots show HA-DA1 and HA-DA1(pep) protein levels. The approximate location of DA1 cleavage sites (arrowed) are shown in UBP15, TCP15 and TCP22. (C) A model of the proposed transient mechanism of DA1 peptidase activation and the consequences of DA1-mediated cleavage of growth regulators during organ growth. PRT1 activity is shown as degrading BB.

## DISCUSSION

### DA1 is a latent peptidase activated by multiple ubiquitylation

The addition of ubiquitin molecules to substrate proteins is a common post-translational modification with many regulatory roles (Hoeller and Dikic 2010; Haglund and Stenmark 2006). We showed that DA1 was ubiquitylated at four lysine residues (Figures 2A-2D, S2) by the E3 ligases BB and DA2, which genetically (Fig. 1A,B) and physically (Figures 1D,E) interact with DA1. Similar levels of DA1 ubiquitylation were observed *in vivo* (Fig. 2I). Loss of function mutants of *BB* and *DA2* synergistically increased the large organ phenotypes of a *DA1* loss of function mutant (Fig. 1A,B) (Li et al. 2008; Xia et al. 2013). Therefore ubiquitylation mediated by these two E3 ligases may increase the growth-limiting activity of DA1. Quantitative growth measurements of leaves (Fig. 1C) showed that individually the *da1-1* mutant and *bb-eod1-2* mutants had distinctive effects on leaf growth; *da1-1* had a delayed time of maximum growth rate, while *bb-eod1-2* showed a slightly accelerated time of maximum growth, compared to Col-0. In combination the mutants showed an even greater maximum growth rate and delayed time of maximum growth rate, increasing final organ size. BB and DA2 act independently to influence final organ size (Disch et al. 2006b), but they both also interact with DA1 (Li et al. 2008; Xia et al. 2013) (Fig. 1A,B), indicating that BB and DA2 may ubiquitylate different substrates, perhaps for proteasomal-mediated degradation. Quantitatively, loss of function mutations in *BB* and *DA2* show their independent effects on growth are less than that of *da1-1* (Fig. 1A,B).

Members of the DA1 family have a canonical MA clan zinc metallopeptidase active site domain in their conserved C terminal region (Supplemental Fig. S3) (Rawlings et al. 2012; Tholander et al. 2010) that was required for limiting growth (Fig. 3F). BB- and DA2-mediated ubiquitylation activated DA1 peptidase activity *in vitro* and in transiently expressed protoplasts (Fig. 3A,E), establishing a biochemical foundation for their joint activities in growth control. Zinc metallopeptidases are maintained in an inactive form by a “cysteine switch” (Van Wart and Birkedal-Hansen 1990) that coordinates a cysteine residue with the zinc atom at the active site to block it. Conformational changes release this and activate the peptidase.

Ubiquitylation of DA1 has the potential to trigger a conformational change that may release inhibition of peptidase activity. (Hoeller et al. 2006) showed that UBD- and UIM-mediated monoubiquitylation of endocytotic proteins including epsin led to a conformational change, mediated by intramolecular interactions between UBDs/UIMs and *cis*-ubiquitin, which regulated endocytosis. The binding of ubiquitin to DA1 UIMs was required for DA1 function *in vivo* (Fig. 2H), and the UIMs conferred similar patterns of ubiquitylation on the heterologous protein GST as seen for DA1 (Fig. 2G for DA1 and Fig. 2E,F for GST-UIM1+2). Related observations were seen in the monoubiquitylation of epsin (Oldham et al. 2002) through coupled monoubiquitylation (Woelk et al. 2006), where UIMs recruit the UIM-containing protein to the ubiquitylation machinery by direct interaction with ubiquitin coupled to ubiquitin donor proteins (Haglund and Stenmark 2006). Mutation of cysteine 274 in the C-terminal zinc finger loop of the LIM-like domain of DA1 abrogated both ubiquitylation and peptidase activity (Fig. 6A,B), suggesting a functional role for this ancient conserved LIM-like domain (Burroughs et al. 2011) (Supplemental Fig. S7) in peptidase activation. Analyses of conformational changes caused by DA1 ubiquitylation and their influence on peptidase activity are required to establish this potential mechanism.

### DA1 cleavage destabilizes its activating E3 ligases BB and DA2, and cleavage of BB leads to targeting by the N-recognin PRT1

The RING E3 ligases BB and DA2 activate DA1 peptidase by ubiquitylation, and are also cleaved by DA1 peptidase (Fig. 3A,B,E,G and Fig. 4). Once cleaved, DA2 appeared to be destabilized in transiently expressed protoplasts (Fig. 3E). Identification of the DA1 cleavage site in BB (Fig. 5A,B) revealed Y61-BB at the neo-N terminus of cleaved BB. This neo-N terminus conferred proteasome-mediated degradation in a cell free system (Fig. 5D). This degradation depended on recognition of the neo-N terminus by the Arabidopsis E3 ligase PRT1 (Fig. 5E,F) (Potuschak et al. 1998; Stary 2003), an N-recognin catalyzing N-end rule mediated degradation (Varshavsky 2011) with a suggested preference for aromatic amino acid N-termini. Interestingly, the neo-N terminal MY61-BB, which was used to express a cleaved version of BB *in planta*, conferred strong proteasome-independent instability (Fig. 5D), in a mechanism that is not yet clear. The lack of MY61-BB function *in vivo* (Fig. 5C) supported the observation that DA1-mediated cleavage of BB leads to its loss of function *in vivo*. Over-expression of *BB* strongly reduced growth, as expected from its inhibitory role in growth (Disch et al. 2006a). The reversal of this inhibition by over-expression of *DA1*, which reduced RFP-BB levels (Fig. 5H) and reversed growth inhibition (Fig. 5G), is consistent with a mechanism involving DA1-mediated reduction of BB activity *via* peptidase-mediated cleavage and subsequent degradation by the N-end rule pathway. Such an activation-destruction mechanism mediated by BB, DA2 and DA1 may provide a way of tightly controlling peptidase activity. These types of mechanisms, often involving ubiquitylation and proteolytic degradation, drive uni-directional cellular processes, for example in cell cycle progression (Reed 2003).

### DA1 peptidase activity also cleaves diverse growth regulators

We previously showed that *TCP14* and *TCP15* function downstream of DA1 and other family members in controlling organ size in Arabidopsis, and reduced function of DA1 family members led to increased TCP14 and TCP15 protein levels (Peng et al. 2015). Similarly, levels of UBP15 protein, which promotes cell proliferation (Lui et al., 2008) and also functions downstream of DA1, were increased in the *da1-1* reduced function mutant (Du et al., 2014). We showed that TCP15 and the related TCP22, and UBP15, were cleaved by DA1 peptidase activity (Fig. 7A,B), but we could not reliably detect TCP14 cleavage by DA1 or DAR1. DA1-mediated cleavage of TCP15 and UBP15 is a plausible mechanism that accounts for these observed reduced protein levels, similar to DA1-mediated inactivation and destabilization of BB by peptidase cleavage. Taken together, these observations suggest a mechanism (Fig. 7C) in which DA1 peptidase, activated transiently by BB or DA2, coordinates a “one-way” cessation of cell proliferation and the initiation of endoreduplication through the cleavage and potential inactivation of proteins that promote cell proliferation and inhibit endoreduplication.

## EXPERIMENTAL PROCEDURES

### Plant Materials, Growth Conditions, and Organ Size Measurements

Arabidopsis thaliana Columbia (Col-0) was the wild-type plant used. Plants were grown in growth-rooms at 20°C with 16 h day/8 h dark cycles, using either soil or MS medium supplemented with 0.5% glucose. Petal and seed areas were imaged by high resolution scanning (3600dpi: Hewlett Packard Scanjet 4370) and analyzed using ImageJ software (http://rsbweb.nih.gov/ij/).

### *In vitro* DA1-mediated Cleavage Assays

FLAG-DA1 was ubiquitylated *in vitro* using either DA2-HIS or BB-HIS as E3 ligases, purified using FLAG-magnetic beads, quantified, and 100 ng added to 100ng BB-HIS, DA2-HIS or BBR-HIS in a 30 µl reaction in 50mM Tris HCl pH 7.4, 5 mM MgCl_2_. Reactions were carried out at 30°C for 4 h and terminated by the addition of SDS sample buffer.

### Mass Spectrometry Analysis

DA1 ubiquitylation patterns were determined from trypsinized proteins purified on SDS-PAGE gels. For LC-MS/MS analysis, peptides were applied to an LTQ-Orbitrab^TM^ (Thermo-Fischer, Waltham, MA, USA) using a nanoAcquity^TM^ UPLC system (Waters Ltd, Manchester, UK). Further details are provided in Supplemental Experimental Procedures.

## AUTHOR CONTRIBUTIONS

M.W.B., J.D., H.D., F.L, H.V., N.D., Y.L. and D.I. designed the research, H.D., F.L., J.D., R.P., H.V., C.N., M.K., C.S., N.McK., L.N., H.V., T.X., L.C., G.S. N.G. and M.S. performed the research and analyzed the data, and M.W.B. wrote the paper.

## ACKNOWLEDGEMENTS

We thank Dr Paul Thomas (Henry Wellcome Laboratory for Cell Imaging, University of East Anglia) for advice and operating the multiphoton microscope, and Dr Cristoph Bücherl (The Sainsbury Laboratory, Norwich) for advice on FRET. We thank Dr Ross Carter (John Innes Centre) for fitting the graph in Figure 1C. We thank Shimadzu Europa GMBH (Duisberg, Gemany) for carrying out Edman sequencing. We thank Andreas Bachmair for the *prt1* EMS allele, and Yukiko Yoshida for the TR-TUBE construct. This work was supported by the Biological and Biotechnological Sciences Research Council (BBSRC) Grant BB/K017225 and Strategic Programme Grant BB/J004588 to MB, and European Commission Contract 037704 (AGROnomics) to MB and DI. JD was supported by a BBSRC CASE Studentship, CN was supported by a PhD Fellowship from from the Landesgraduiertenförderung Sachsen-Anhalt, and ND by an Independent Junior Research Group grant from the ScienceCampus Halle - Plant-based Bioeconomy, the Deutsche Forschungsgemeinschaft (DFG; grant DI 1794/3-1), the DFG Graduate Training Centre GRK1026 and the Leibniz Institute of Plant Biochemistry. YL was supported by the National Natural Science Foundation of China (Grants 91417304; 31425004; 91017014; 31221063; 31100865), National Basic Research Program of China (Grant 2009CB941503) and the Ministry of Agriculture of China (Grant 2013ZX08009-003-003). MB and YL are in the CASJIC Centre of Excellence in Plant and Microbial Sciences (CEPAMS).

## REFERENCES

Andriankaja M, Dhondt S, De Bodt S, Vanhaeren H, Coppens F, De Milde L, Mühlenbock P, Skirycz A, Gonzalez N, Beemster GTS, et al. 2012. Exit from Proliferation during Leaf Developmentin Arabidopsis thaliana: A Not-So-Gradual Process. Developmental Cell 22: 64–78.

Bachmair A, Finley D, Varshavsky A. 1986. In vivo half-life of a protein is a function of its amino-terminal residue. Science 234: 179–186.

Barry ER, Camargo FD. 2013. The Hippo superhighway: signaling crossroads converging on the Hippo/Yap pathway in stem cells and development. Curr Opin Cell Biol 25: 247–253.

Breuer C, Ishida T, Sugimoto K. 2010. Developmental control of endocycles and cell growth in plants. Current Opinion in Plant Biology 13: 654–660.

Breuninger H, Lenhard M. 2012. Expression of the central growth regulator BIG BROTHER is regulated by multiple *cis*-elements. BMC Plant Biology 12: 41.

Burroughs AM, Iyer LM, Aravind L. 2011. Functional diversification of the RING finger and other binuclear treble clef domains in prokaryotes and the early evolution of the ubiquitin system. Mol BioSyst 7: 2261.

De Veylder L, Larkin JC, Schnittger A. 2011. Molecular control and function of endoreplication in development and physiology. Trends in Plant Science 16: 624–634.

Disch S, Anastasiou E, Sharma VK, Laux T, Fletcher JC, Lenhard M. 2006a. The E3 Ubiquitin Ligase BIG BROTHER Controls Arabidopsis Organ Size in a Dosage-Dependent Manner. Current Biology 16: 272–279.

Du L, Li N, Chen L, Xu Y, Li Y, Zhang Y, Li C, Li Y. 2014. The Ubiquitin Receptor DA1 Regulates Seed and Organ Size by Modulating the Stability of the Ubiquitin-Specific Protease UBP15/SOD2 in Arabidopsis. The Plant Cell 26: 665–677.

Efroni I, Blum E, Goldshmidt A, Eshed Y. 2008. A protracted and dynamic maturation schedule underlies Arabidopsis leaf development. The Plant Cell 20: 2293–2306.

Faden F, Ramezani T, Mielke S, Almudi I, Nairz K, Froehlich MS, Höckendorff J, Brandt W, Hoehenwarter W, Dohmen RJ, et al. 2016. Phenotypes on demand via switchable target protein degradation in multicellular organisms. Nature Communications 7: 12202.

Green AA, Kennaway JR, Hanna AI, Bangham JA, Coen E. 2010. Genetic control of organ shape and tissue polarity. PLoS Biol 8: e1000537.

Haglund K, Di Fiore PP, Dikic I. 2003. Distinct monoubiquitin signals in receptor endocytosis. Trends in Biochemical Sciences 28: 598–603.

Haglund K, Stenmark H. 2006. Working out coupled monoubiquitination. Nature Cell Biology 8: 1218–1219.

Hicke L, Schubert HL, Hill CP. 2005. Ubiquitin-binding domains. Nat Rev Mol Cell Biol 6: 610–621.

Hoeller D, Crosetto N, Blagoev B, Raiborg C, Tikkanen R, Wagner S, Kowanetz K, Breitling R, Mann M, Stenmark H, et al. 2006. Regulation of ubiquitin-binding proteins by monoubiquitination. Nature Cell Biology 8: 163–169.

Hoeller D, Dikic I. 2010. Regulation of ubiquitin receptors by coupled monoubiquitination. Subcell Biochem 54: 31–40.

Husnjak K, Dikic I. 2012. Ubiquitin-Binding Proteins: Decoders of Ubiquitin-Mediated Cellular Functions. Annu Rev Biochem 81: 291–322.

Johnston LA, Gallant P. 2002. Control of growth and organ size inDrosophila. Bioessays 24: 54–64.

Kadrmas JL, Beckerle MC. 2004. The LIM domain: from the cytoskeleton to the nucleus. Nat Rev Mol Cell Biol 5: 920–931.

Kazama T, Ichihashi Y, Murata S, Tsukaya H. 2010. The mechanism of cell cycle arrest front progression explained by a KLUH/CYP78A5-dependent mobile growth factor in developing leaves of Arabidopsis thaliana. Plant and Cell Physiology 51: 1046–1054.

Kim H, Chen J, Yu X. 2007. Ubiquitin-binding protein RAP80 mediates BRCA1-dependent DNA damage response. Science 316: 1202–1205.

Komander D, Rape M. 2012. The Ubiquitin Code. Annu Rev Biochem 81: 203–229.

Li Y, Zheng L, Corke F, Smith C, Bevan MW. 2008. Control of final seed and organ size by the DA1 gene family in Arabidopsis thaliana. Genes & Development 22: 1331–1336.

Oldham CE, Mohney RP, Miller SLH, Hanes RN, O'Bryan JP. 2002. The ubiquitin-interacting motifs target the endocytic adaptor protein epsin for ubiquitination. Current Biology 12: 1112–1116.

Pan D. 2010. The hippo signaling pathway in development and cancer. Developmental Cell 19: 491–505.

Peng Y, Chen L, Lu Y, Wu Y, Dumenil J, Zhu Z, Bevan MW, Li Y. 2015. The Ubiquitin Receptors DA1, DAR1, and DAR2 Redundantly Regulate Endoreduplication by Modulating the Stability of TCP14/15 in Arabidopsis. The Plant Cell 27: 649–662.

Potuschak T, Stary S, Schlögelhofer P, Becker F, Nejinskaia V, Bachmair A. 1998. PRT1 of Arabidopsis thaliana encodes a component of the plant N-end rule pathway. Proc Natl Acad Sci USA 95: 7904–7908.

Rawlings ND, Barrett AJ, Bateman A. 2012. MEROPS: the database of proteolytic enzymes, their substrates and inhibitors. Nucleic Acids Research. 40, D343–D350.

Reed SI. 2003. Ratchets and clocks: the cell cycle, ubiquitylation and protein turnover. Nat Rev Mol Cell Biol 4: 855–864.

Sato Y, Yoshikawa A, Mimura H, Yamashita M, Yamagata A, Fukai S. 2009. Structural basis for specific recognition of Lys 63-linked polyubiquitin chains by tandem UIMs of RAP80. The EMBO Journal 28: 2461–2468.

Sluis A, Hake S. 2015. Organogenesis in plants: initiation and elaboration of leaves. Trends Genet 31: 300–306.

Stary S. 2003. PRT1 of Arabidopsis Is a Ubiquitin Protein Ligase of the Plant N-End Rule Pathway with Specificity for Aromatic Amino-Terminal Residues. Plant Physiology 133: 1360–1366.

Tholander F, Roques B-P, Fournié-Zaluski M-C, Thunnissen MMGM, Haeggström JZ. 2010. Crystal structure of leukotriene A4 hydrolase in complex with kelatorphan, implications for design of zinc metallopeptidase inhibitors. FEBS Letters 584: 3446–3451.

van der Krogt GNM, Ogink J, Ponsioen B, Jalink K. 2008. A Comparison of Donor-Acceptor Pairs for Genetically Encoded FRET Sensors: Application to the Epac cAMP Sensor as an Example ed. K.-W. Koch. PLoS ONE 3: e1916.

Van Wart HE, Birkedal-Hansen H. 1990. The cysteine switch: a principle of regulation of metalloproteinase activity with potential applicability to the entire matrix metalloproteinase gene family. Proc Natl Acad Sci USA 87: 5578–5582.

Varshavsky A. 2011. The N-end rule pathway and regulation by proteolysis. Protein Sci. 20: 1298–1254

Woelk T, Oldrini B, Maspero E, Confalonieri S, Cavallaro E, Di Fiore PP, Polo S. 2006. Molecular mechanisms of coupled monoubiquitination. Nature Cell Biology 8: 1246–1254.

Xia T, Li N, Dumenil J, Li J, Kamenski A, Bevan MW, Gao F, Li Y. 2013. The ubiquitin receptor DA1 interacts with the E3 ubiquitin ligase DA2 to regulate seed and organ size in Arabidopsis. The Plant Cell 25: 3347–3359.

Xu F, Zhu C, Çevik V, Johnson K, Liu Y, Sohn K, Jones JD, Holub EB, Li X. 2015. Autoimmunity conferred by chs3–2D relies on CSA1, its adjacent TNL-encoding neighbour. Sci Rep 5: 8792.

